# Tech Note: Simplified protocol for SMARTer Pico kit

**DOI:** 10.1101/2025.09.02.673416

**Authors:** Orlando Contreras-López, Laia Masvidal, Jun Wang, Elísabet Einarsdóttir

## Abstract

The [SMARTer® Stranded Total RNA-Seq Kit v2 – Pico Input Mammalian](https://www.takarabio.com/) kit from Takara® (SMARTer Pico) has proved successful and reliable in generating stranded RNA Illumina libraries from degraded total RNA and ultra-low input amounts of total RNA below detection level. Here we attempted to streamline and simplify the library prep protocol at the key fragmentation step bottleneck. Our key findings were that reduced fragmentation times neither affect the depletion efficiency, nor the library complexity. Skipping the fragmentation resulted in longer libraries when examined in the capillary electrophoresis but this was compensated for during sequencing, as long fragments are less likely to form clusters during sequencing. Skipping the fragmentation also affected the gene body coverage, where a bias to the 5’ end was observed though this compromised neither the data quality, complexity nor reproducibility. Additionally, using 16 PCR cycles seems to have little effect on the library complexity. Overall, we can see that sample input is the key to library complexity and reproducibility, while fragmentation time has less impact on data.

Is total RNA fragmentation needed? Is rRNA depletion affected by fragmentation time? Or affected by input? Can we get reliable data when starting with low input?

## Introduction

The [SMARTer® Stranded Total RNA-Seq Kit v2 – Pico Input Mammalian](https://www.takarabio.com/) kit from Takara® (SMARTer Pico) has proved successful and reliable in generating stranded RNA Illumina libraries from degraded total RNA and ultra-low input amounts of total RNA below detection level. This kit uses random priming cDNA synthesis of both coding and non-coding RNA which is ideal for processing degraded total RNA input, such as those obtained from formalin-fixed or paraffin-embedded (FFPE) samples.

Moreover, the kit incorporates a technology that enables the removal of mammalian ribosomal cDNA, without loss of other cDNA molecules originating from non-coding or coding RNAs.

The increased demand for low input protocols such as the SMARTer Pico motivated us to look at ways to optimize the processing of samples. In order to accomplish this, we have investigated ways to streamline and simplify the SMARTer Pico library prep protocol (see *Figure 1*). The original protocol contains a fragmentation step, just before the first-strand synthesis, that has to be customized depending on the quality ([RIN value](https://www.agilent.com/cs/library/applications/5989-1165EN.pdf) or [DV200](https://emea.illumina.com/content/dam/illumina-marketing/documents/products/technotes/evaluating-rna-quality-from-ffpe-samples-technical-note-470-2014-001.pdf)) of the total RNA sample. If a sample is of good quality (RIN > 7), the recommended fragmentation time is 4 min; while if the quality is low (RIN < 3), it is recommended to skip the fragmentation. In a given project, our users might have samples with a range of RIN values or it might be impossible to determine the RIN value due to low sample concentration. Therefore, the fragmentation step presents a bottleneck in the library prep since samples within a project may require different treatment.

**Figure 1.**
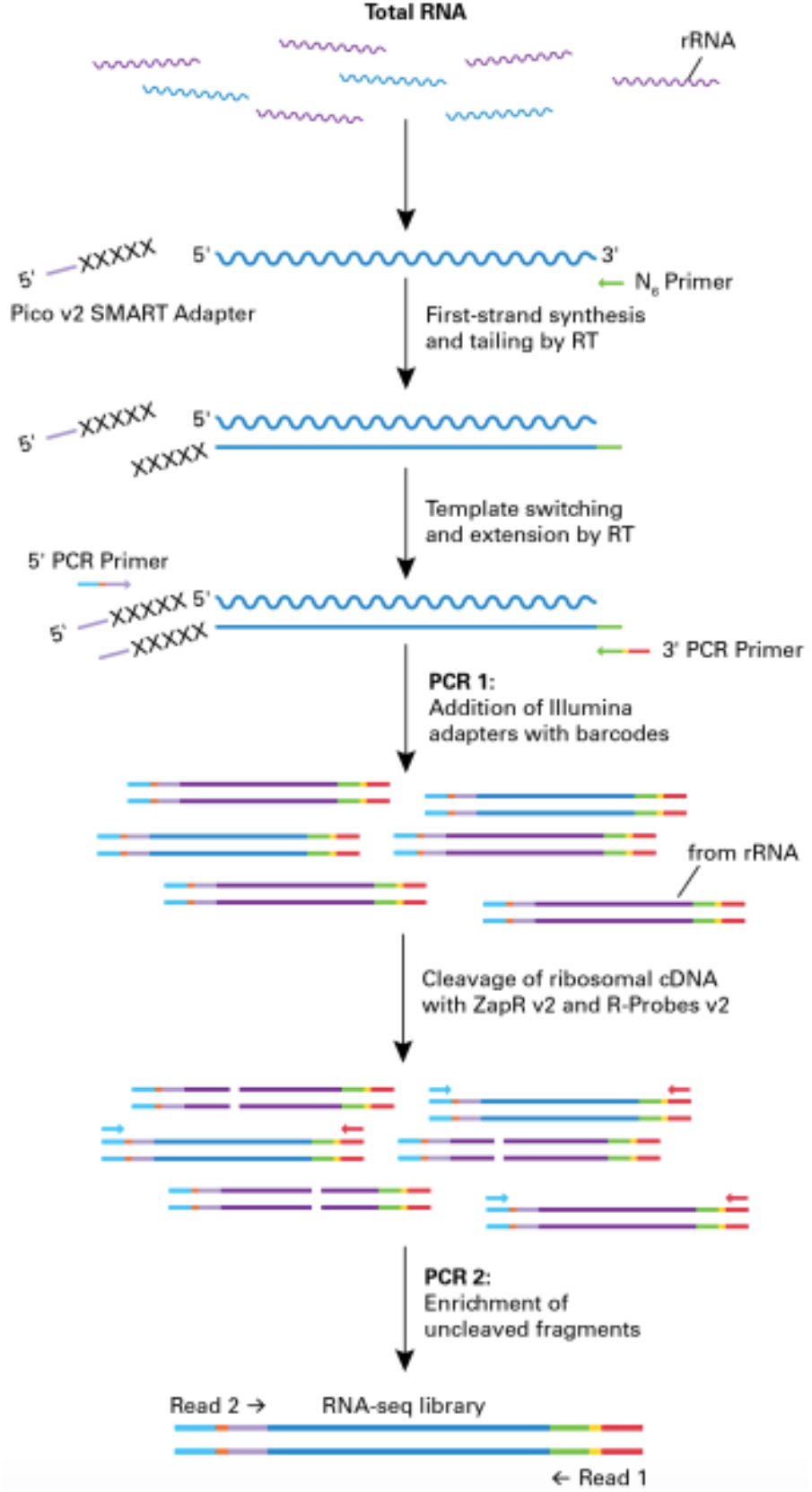
Schematic of the SMARTer Stranded Total RNA-Seq Kit v2 – Pico Input Mammalian protocol. Random priming allows the generation of cDNA from all RNA fragments in the sample, including rRNA and degraded mRNA. The Reverse Transcriptase adds additional nucleotides when it reaches the 5′ end. Those extra nucleotides are used for the synthesis of the second strand of the cDNA. The next step is a short round of PCR amplification which adds full-length Illumina adapters, including barcodes. The cDNA originating from rRNA is then cleaved by ZapR enzyme in the presence of mammalian-specific R-Probes. This process leaves the library with only fragments from non-rRNA molecules. These fragments are enriched via a second round of PCR amplification. Image taken from Takara®.

To investigate the effect of fragmentation time, we prepared libraries using aliquots of a good quality total RNA sample with different fragmentation times. In addition to the fragmentation time, we also included different input amounts (from 1 ng to 10 ng), all at a fixed number of PCR cycles for PCR2 (*Figure 1*). Data from a user project as well as data from Takara were also used for the analysis. The sequencing data allowed us to compare the different conditions, demonstrating that the fragmentation time is not a determinant in the quality of the generated data.

## Methodology

Three different datasets were used to investigate the effects of fragmentation and input amount on the library preparation efficiency and data quality.

**Dataset 1** was generated using Universal Human Reference RNA 2×200 ug from AH diagnostics (cat# 740000). This total RNA was diluted to 1.25 ng/ul and 0.125 ng/ul for the 10 ng and 1ng input, respectively. The total RNA samples were run on the Fragment Analyzer HS RNA, where the 0.125 ng/ul samples were undetectable. The 1.25 ng/ul samples had a RIN 9, confirming the high quality of the samples. Library preparation was carried out in duplicates, using 1 ng or 10 ng as input. The samples were subjected to three fragmentation times: 0, 2, and 4 mins (see *Table 1*). The number of PCR cycles for PCR2 (*Figure 1*) was 16. As expected, the library preparation yields are proportional to the input. Also, the average sizes are larger with shorter fragmentation times (see *Table 1* and *Figure 2*). Samples were sequenced on an Illumina NovaSeq 6000 S4 flowcell, with an average of 200M 2×150 bp reads per sample.

**Table 1.** Library QC.Concentration measurements and average size determination for each library prepared in house starting from Invitrogen™ AM7852 total RNA. QC pass threshold is concentration > 2 nM. All libraries passed QC. Library yield correlates with sample input.

| Library | Concentration (nM) | Average Size (bp) |
| --- | --- | --- |
| 1ng_Frag_0min_R1 | 6.67 | 871 |
| 1ng_Frag_0min_R2 | 5.47 | 864 |
| 10ng_Frag_0min_R1 | 115.44 | 784 |
| 10ng_Frag_0min_R2 | 112.97 | 814 |
| 1ng_Frag_2min_R1 | 12.71 | 418 |
| 1ng_Frag_2min_R2 | 15.48 | 415 |
| 10ng_Frag_2min_R2 | 127.57 | 580 |
| 10ng_Frag_2min_R1 | 127.82 | 618 |
| 1ng_Frag_4min_R1 | 20.61 | 328 |
| 1ng_Frag_4min_R2 | 19.16 | 334 |
| 10ng_Frag_4min_R1 | 120.68 | 501 |
| 10ng_Frag_4min_R2 | 102.78 | 582 |

**Figure 2.**
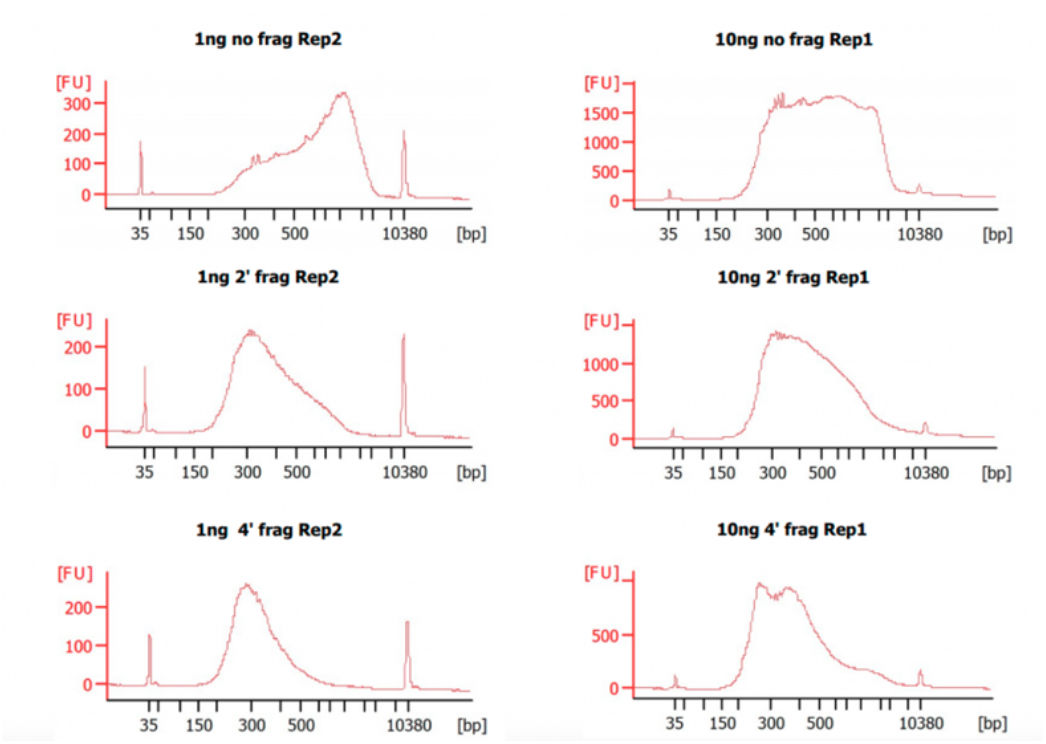
Bioanalyzer electrpherograms of the final libraries. Bioanalyzer High Sensitivity DNA Assay was used to assess the size distribution of the final libraries from Dataset 1. Samples without fragmentation show a higher proportion of fragments above 1000 bp.

**Dataset 2** was generated using a human total RNA sample provided by an NGI user. Three different input amounts were used: 3, 10, and 30 ng. The RIN value determined with the Fragment Analyzer HS RNA of these samples was >9.4. The samples were fragmented for 4 mins and 11 cycles were performed for PCR2. Samples were sequenced on an Illumina NovaSeq 6000 S4, 2×150 bp reads, with an average of 200M reads per sample. Samples were named *Unsub* (*Figures 7-9*).

**Dataset 3** was obtained from Takara’s [TechNote on SMARTer Pico](https://www.takarabio.com/learning-centers/next-generation-sequencing/technical-notes/rna-seq/stranded-libraries-from-picogram-input-total-rna-(v2)). Here, Takara used human lung FFPE total RNA without fragmentation, with 16 PCR cycles for PCR2. The dataset had the merged data from two library preps for each input, with 18M and 54.6M reads for the 1 ng and 10 ng input merged data, respectively.

All datasets were analyzed using the [nfcore/rnaseq](https://github.com/nf-core/rnaseq) pipeline. The same software versions and options were used for all the analyses, and MultiQC was used to extract data for all the figures.

## Results

### Sample input, not fragmentation time, determines library similarity

In order to determine the effect of the fragmentation time and/or sample input on the data, the libraries from **Dataset 1** were clustered. The dendrogram is generated from the normalized gene counts through edgeR. Euclidean distances between log2 normalized CPM values are then calculated and clustered. The clustering result shown in *Figure 3* indicates that libraries from samples with 10 ng input are more similar to each other than to those with 1 ng.

**Figure 3.**
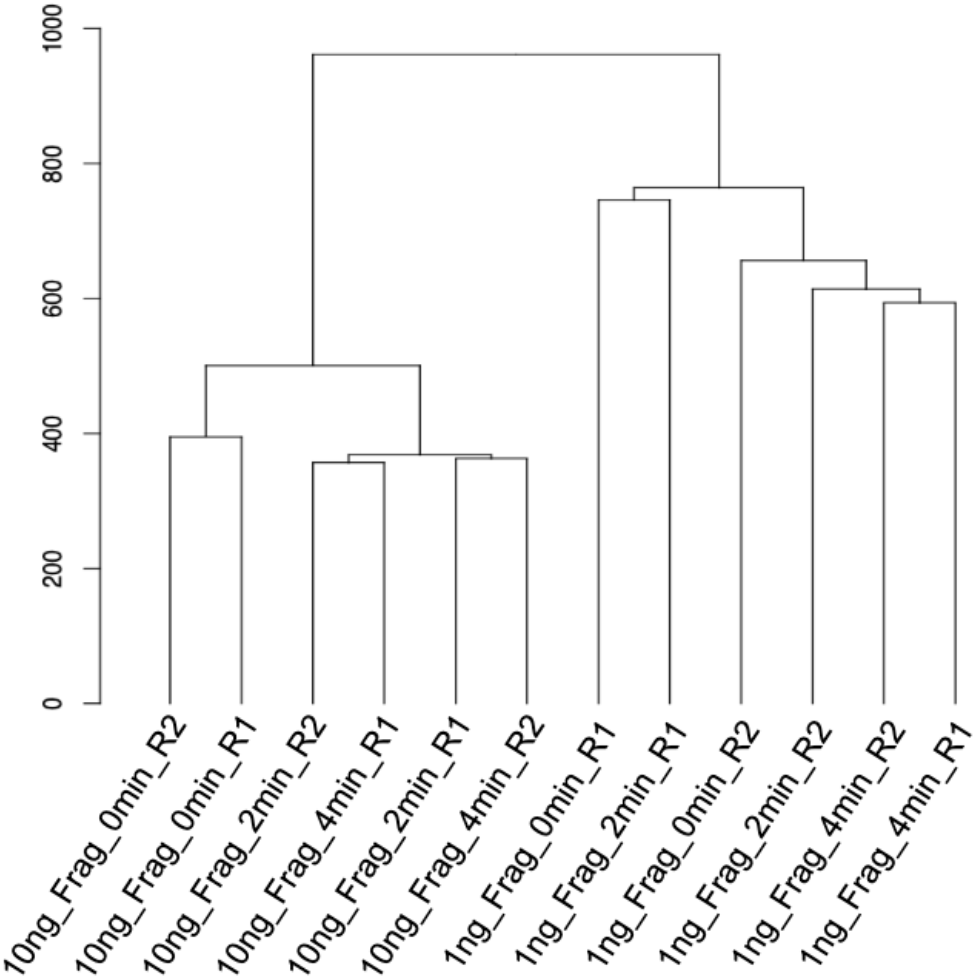
Dendrogram showing sample clustering. Euclidean distances between log2 normalized CPM values are then calculated and clustered. Libraries from Dataset 1 are grouped by input regardless of the fragmentation time used for library prep.

Furthermore, the fragmentation time of 2 or 4 mins has little impact on the library data when 10 ng input is used, as those samples fall within the same clades. Lower input leads to lower reproducibility, regardless of the fragmentation time.

We assessed the percentages of reads aligned to the reference genome, as well as the percentage of reads assigned to features in the genome (*Figure 4*). We found no large differences among the datasets suggesting that regardless of the input and fragmentation time, the proportion of useful reads remains the same.

**Figure 4.**
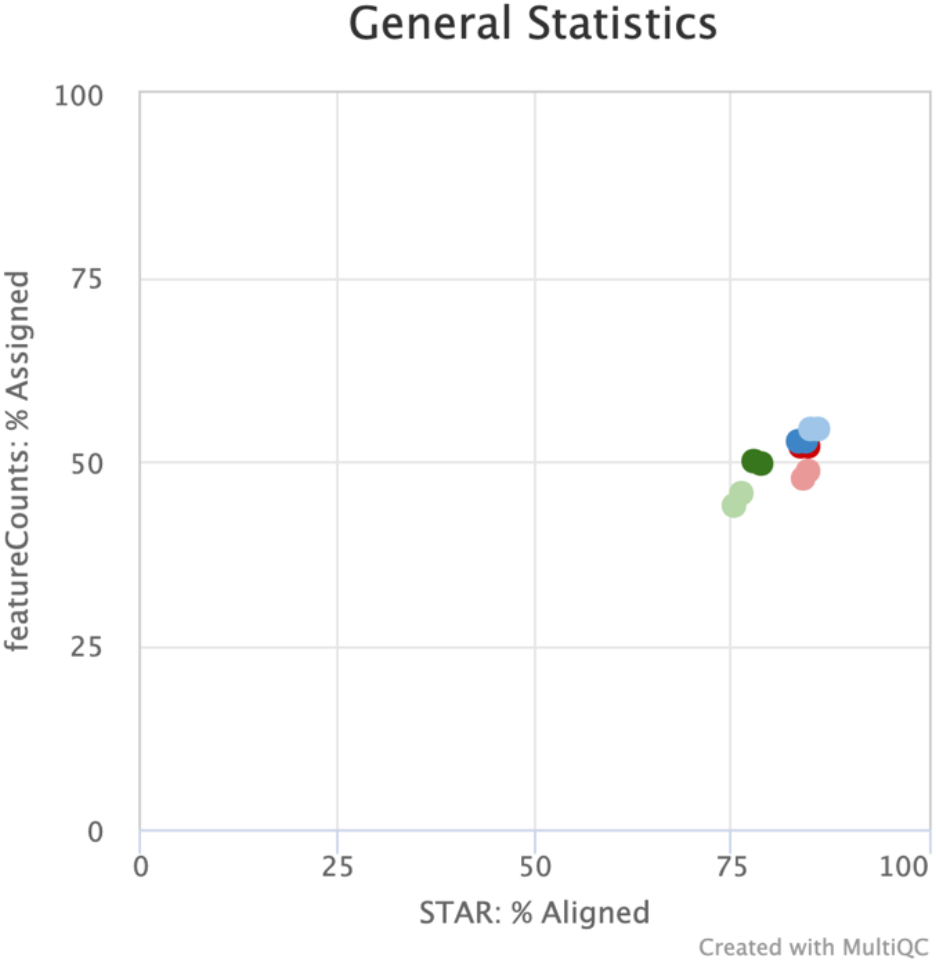
Scatter plot showing read the percentage of reads assigned to a genome feature and percentage of reads aligned to the genome. Data were obtained from the MultiQC report. Percentage of Assigned reads were acquired from [Subread featureCounts](http://bioinf.wehi.edu.au/featureCounts/) that counts mapped reads for genomic features such as genes, exons, promoter, gene bodies, genomic bins, and chromosomal locations. The percentage of aligned reads was determined with [STAR](https://github.com/alexdobin/STAR) which is an ultrafast universal RNA-seq aligner.

time has less impact on data.

### Library insert size is affected by fragmentation time

To test if longer fragmentation times led to shorter library inserts, we estimated the insert size distribution based on the data from the [inner distance](http://rseqc.sourceforge.net/#inner-distance-py) module within the MultiQC report. The distance reported is the mRNA length between two paired reads. To this value, the length of the paired reads was added to get the insert size. *Figure 5* shows the frequency of the insert size for the libraries from **Dataset 1**. The samples subjected to longer fragmentation times (red and green) have shorter inserts, on average. However, this is smaller than the difference in average size of the library obtained from the capillary electrophoresis (see *Table 1*); the difference there was as large as 500 bp compared to 80 bp in the sequencing data. This could be explained by the difference in clustering efficiency between short and long fragments, leading to an underrepresentation of the very long fragments in the sequencing data.

**Figure 5.**
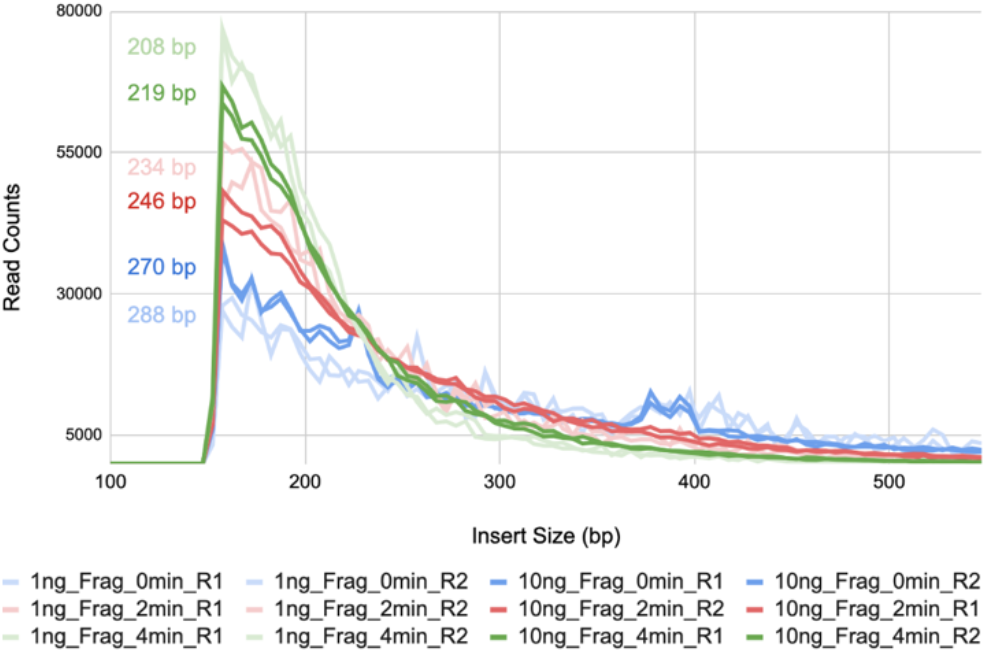
Insert size distribution. Insert size was estimated using the data obtained from the Inner Distance module from [RSeQC](http://rseqc.sourceforge.net/) in the MultiQC report. Average Insert Size is shown for each library in their respective color. Shorter fragmentation times lead to longer libraries.

Overall, the data shows that the fragmentation time does reduce the size of the library inserts.

### Library complexity depends on sample input and is not affected by fragmentation time

Library complexity is a measure of the number of unique molecules in the library. For any application, the goal is to make the libraries as complex as possible. This parameter is helpful in determining where sensitivity loss can occur (detection of low expressed genes) or where sequencing errors are created (duplicates). It is therefore important to determine if the fragmentation time affects library complexity and if so, to what degree.

Library complexity was estimated using the [preseq](http://smithlabresearch.org/software/preseq/) module from the MultiQC report. *Figure 6* shows the rarefaction curves for **Dataset 1** (blue, red and green), **Dataset 2**, the user sample (orange), and **Dataset 3**, from Takara (grey). One can see that the library complexity is more dependent on the sample input, and not so much on the fragmentation time.

**Figure 6.**
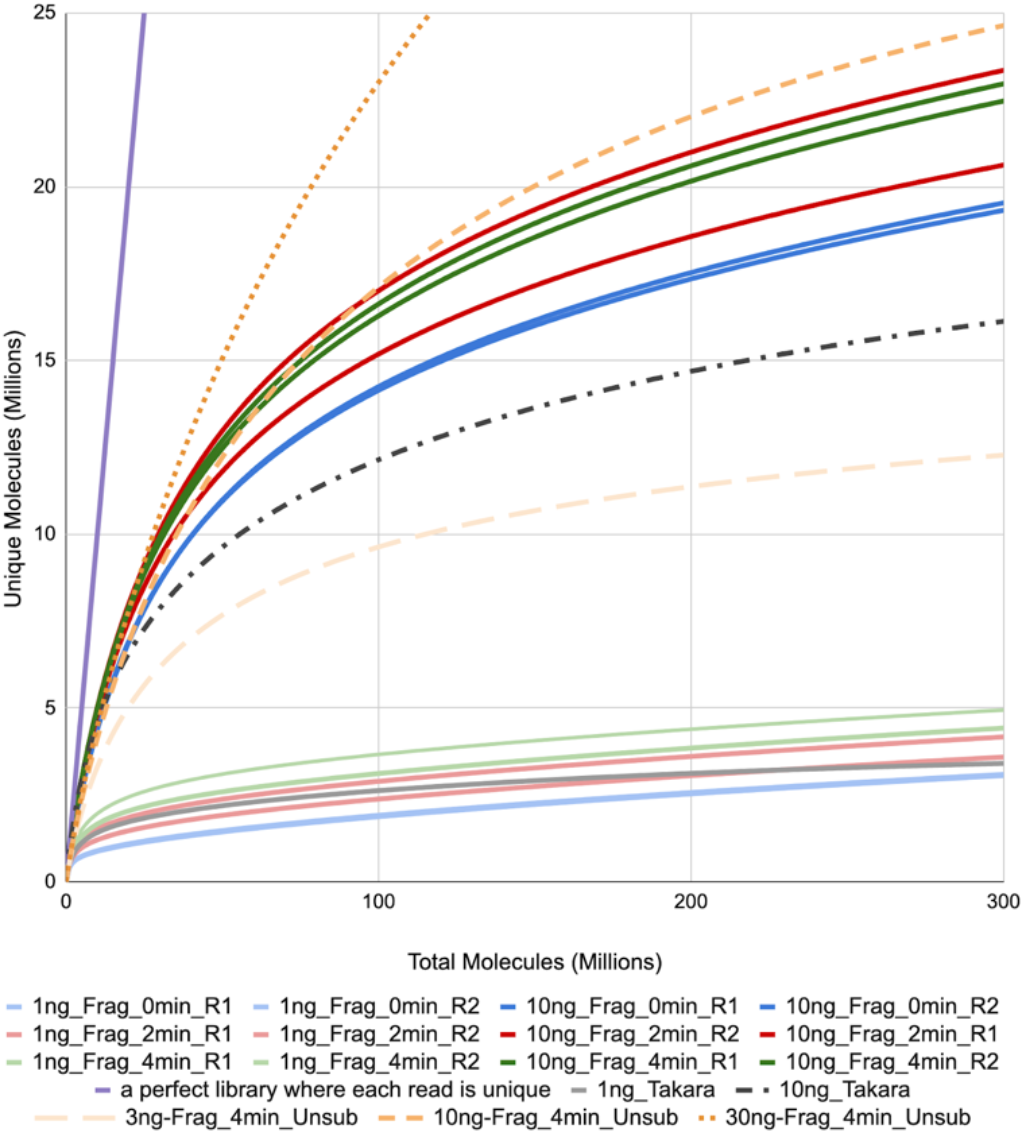
Library Complexity Curve. Complexity was estimated using the data obtained from the Complexity of the MultiQC report. Libraries group by input regardless of the fragmentation time used for library prep. The purple line shows the curve for an ideal library where each molecule is unique.

**Figure 7.**
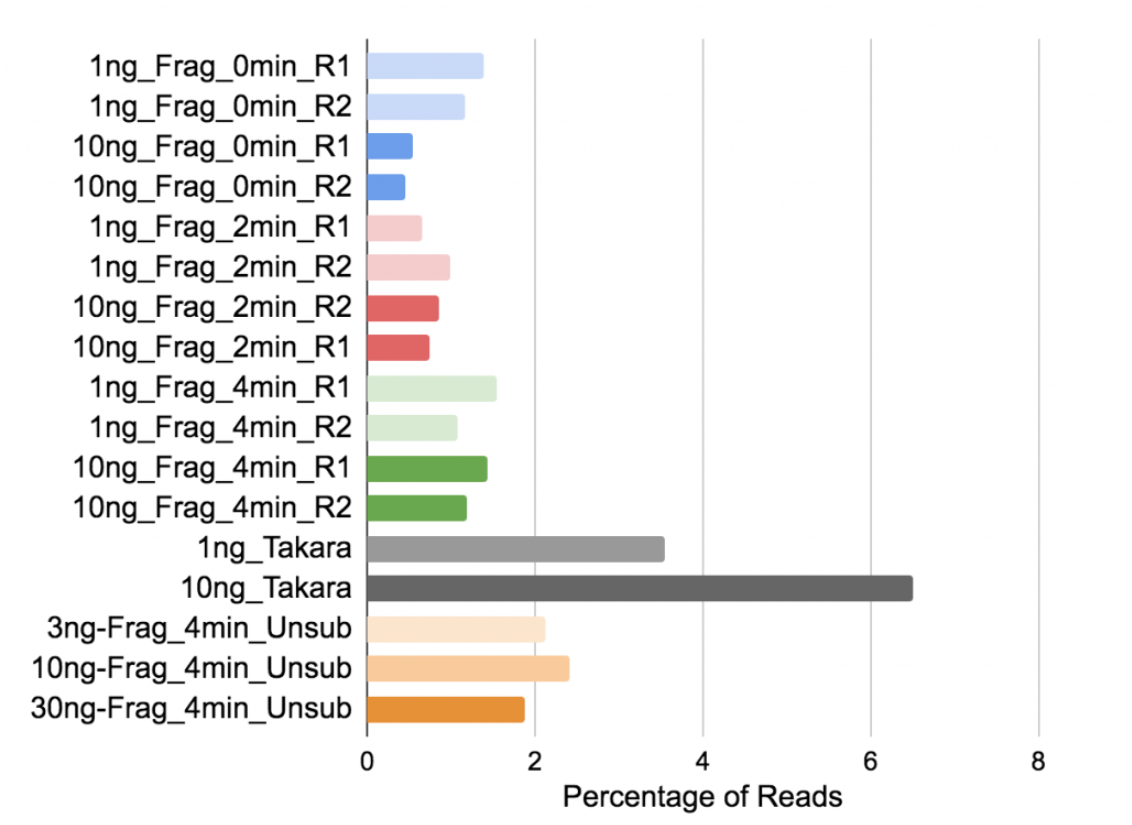
Percentage of reads mapping to ribosomal RNA. Biotype Counts shows reads overlapping genomic features of different biotypes, counted by [featureCounts](http://bioinf.wehi.edu.au/featureCounts/).

### rRNA depletion efficiency is independent of sample input and fragmentation time

One of the key features of the SMARTer Pico kit is the depletion of rRNA in mammalian samples. The removal of the rRNA is dependent on probes that target these sequences and are then enzymatically degraded (*Figure 1*). According to the manufacturer, excessive fragmentation can lead to an inefficient depletion process, but we did not expect a difference in the depletion efficiency when fragmentation was skipped on good quality samples from **Dataset 1** or **Dataset 2**.

The depletion efficiency was estimated from the percentage of reads overlapping the genomic region spanned by the rRNA feature. The higher the percentage of reads mapped to rRNA, the lower the depletion efficiency.

*Figure 7* shows that neither the fragmentation time nor the sample input affect the depletion efficiency. On the other hand, sample quality seems to be crucial since the libraries from **Dataset 3**, generated from FFPE samples, show lower depletion efficiency than the high-quality samples from **Dataset 1 or Dataset 2**.

### Mitochondrial rRNA depletion efficiency is affected neither by sample input nor fragmentation time

Another feature of the SMARTer Pico kit is the depletion of Human Mitochondrial rRNA (Mt rRNA). The removal of the Mt rRNA is also dependent on probes that target these sequences and are then enzymatically degraded (*Figure 1*). Similar to the analysis above, a higher percentage of reads mapping to Mt rRNA indicates lower depletion efficiency. *Figure 8* shows that the depletion of the Mt rRNA is sample dependent (tissue or cell type) and it is neither clearly affected by the fragmentation time nor the sample input.

**Figure 8.**
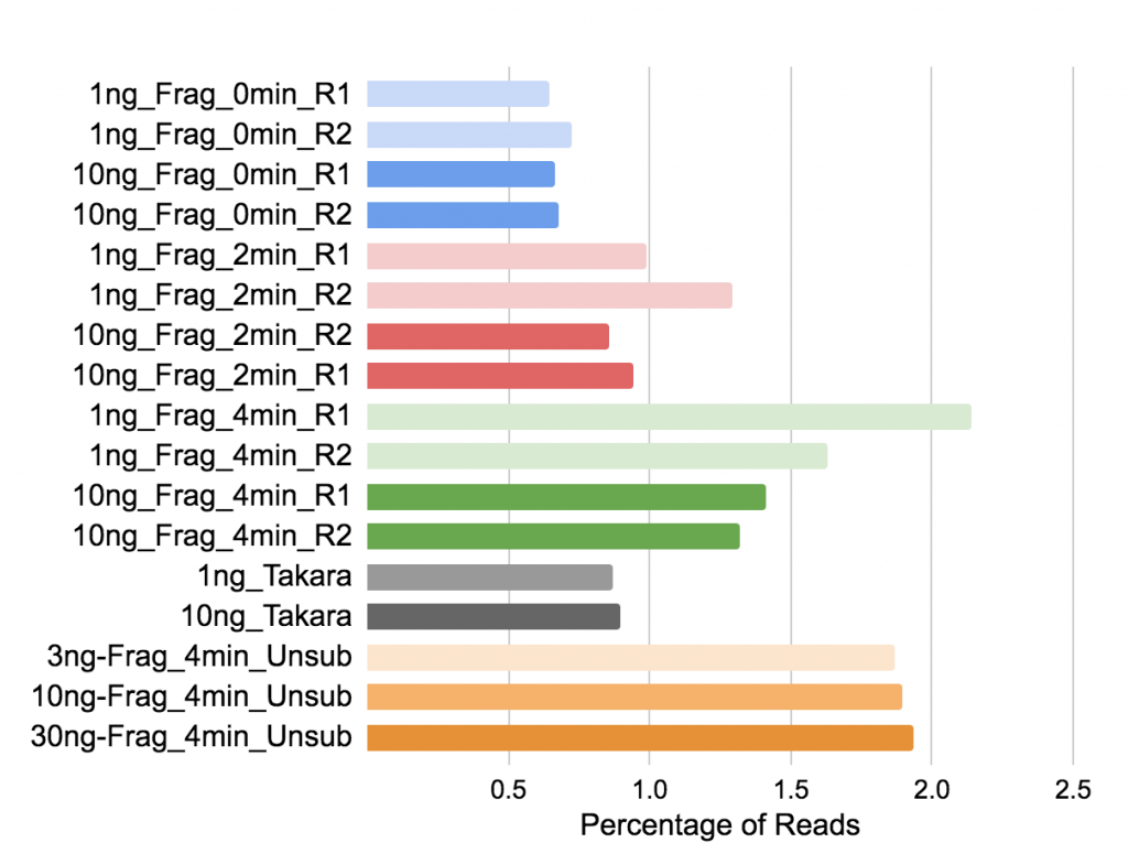
Percentage of reads mapping to mitochondrial ribosomal RNA. Biotype Counts shows reads overlapping Mt_rRNA, counted by [featureCounts](http://bioinf.wehi.edu.au/featureCounts/) in the MultiQC report.

### Gene body coverage is biased towards the 5’ end when fragmentation is not performed

Gene Body Coverage calculates read coverage over gene bodies. This is used to check if read coverage is uniform and if there is any 5′ or 3′ bias. *Figure 9* shows that samples from all datasets have very similar gene body coverage, except for the high-quality samples that were not subjected to fragmentation (blue).

**Figure 9.**
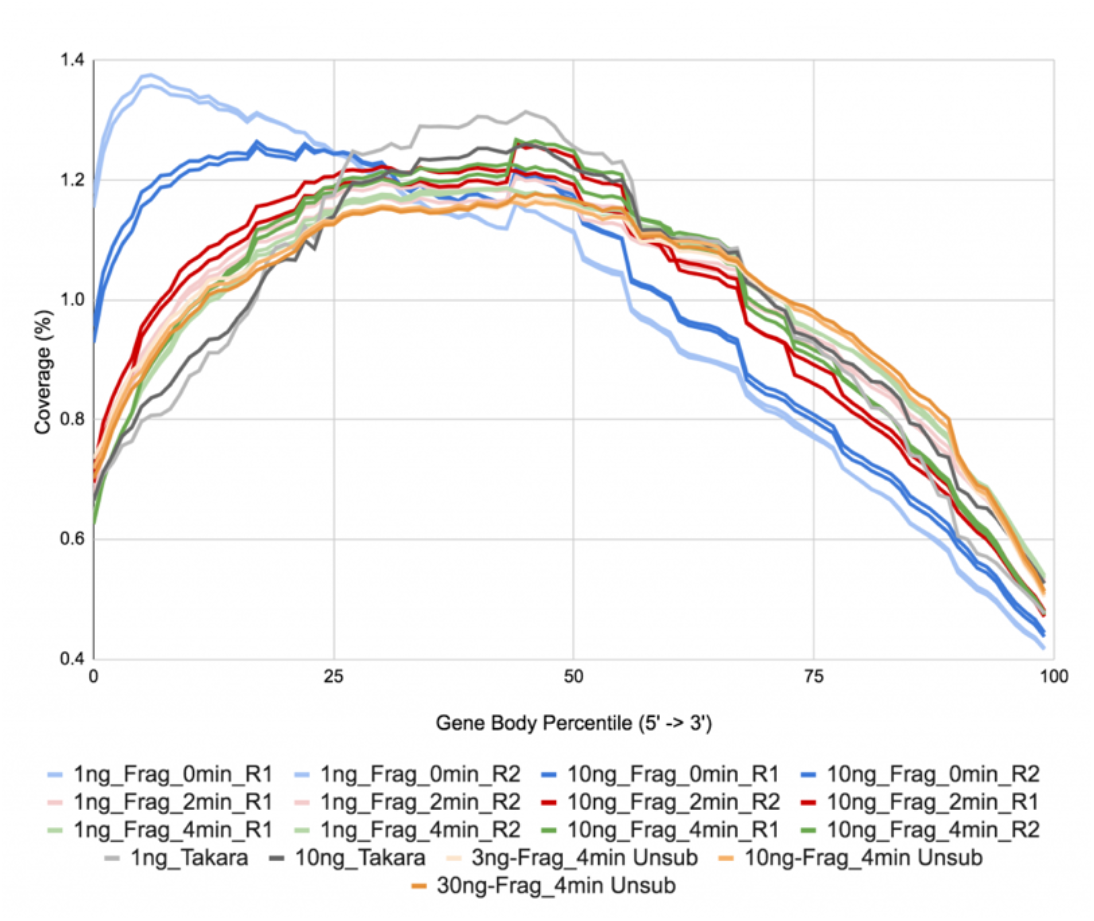
Gene body coverage.Gene Body Coverage is obtained from the module called the same from RSeQC in the MultiQC report.

These samples show a bias to the 5’ end of the gene body but with a similar coverage on the rest of the gene body when compared to the rest of the samples. Other inputs or fragmentation times do not change the gene body coverage.

## Conclusion

In this tech note, we aimed to determine the effects of different fragmentation times and input amounts on the data produced by the SMARTer Pico prep. Although we lack statistical power, this approach can guide us to the following conclusions: The shorter than recommended fragmentation times neither affect the depletion efficiency (*Figures 7* and *8*) nor the library complexity (*Figure 6*).

Skipping the fragmentation resulted in longer libraries when examined in the capillary electrophoresis (*Figure 2*) but this is compensated for during sequencing, as long fragments are less likely to form clusters during sequencing (*Figure 5*). Skipping the fragmentation also affected the gene body coverage, where a bias to the 5’ end was observed (*Figure 9)* but this compromised neither the data quality (*Figure 4*), the complexity (*Figure 6*) nor the reproducibility (*Figure 3*).

Additionally, using 16 PCR cycles seems to have little effect on the library complexity (*Figure 6*). Overall, we can see that sample input is the key to library complexity and reproducibility, while fragmentation time has less impact on data.

